# pOPIN-GG: A resource for modular assembly in protein expression vectors

**DOI:** 10.1101/2021.08.10.455798

**Authors:** Adam R. Bentham, Mark Youles, Melanie N. Mendel, Freya A. Varden, Juan Carlos De la Concepcion, Mark J. Banfield

## Abstract

The ability to recombinantly produce target proteins is essential to many biochemical, structural, and biophysical assays that allow for interrogation of molecular mechanisms behind protein function. Purification and solubility tags are routinely used to maximise the yield and ease of protein expression and purification from *E. coli*. A major hurdle in high-throughput protein expression trials is the cloning required to produce multiple constructs with different solubility tags. Here we report a modification of the well-established pOPIN expression vector suite to be compatible with modular cloning via Type IIS restriction enzymes. This allows users to rapidly generate multiple constructs with any desired tag, introducing modularity in the system and delivering compatibility with other modular cloning vector systems, for example streamlining the process of moving between expression hosts. We demonstrate these constructs maintain the expression capability of the original pOPIN vector suite and can also be used to efficiently express and purify protein complexes, making these vectors an excellent resource for high-throughput protein expression trials.

**Highlights:**

- pOPIN-GG expression vectors allow for modular cloning enabling rapid screening of purification and solubility tags at no loss of expression compared to previous vectors.
- Cloning into the pOPIN-GG vectors can be performed from PCR products or from level 0 vectors containing the required parts.
- Several vectors with different resistances and origins of replication have been generated allowing the effective co-expression and purification of protein complexes.
- All pOPIN-GG vectors generated here are available on Addgene, as well as level 0 acceptors and tags.

## Introduction

Understanding protein function is key to answering many biological questions. Biochemical, structural, and biophysical techniques that probe the molecular mechanisms behind protein function are reliant on the production of purified protein for use in these assays. Procedures for protein expression and purification from *Escherichia coli* have been advanced by methodologies which allow for the high-throughput generation of constructs (Rosano and Ceccarelli, 2014). Intrinsic to generating soluble protein of the more difficult targets in *E. coli* is the capacity to test multiple solubility tags, such as the Small Ubiquitin-like modifier (SUMO) or the Maltose Binding Protein (MBP) tags, which allow for the production of proteins that would be otherwise insoluble (di Guana et al., 1988; Malakhov et al., 2004). Further, the use of purification tags frequently allows for rapid capture of proteins of interest from cell lysates. However, purification and solubility tags are often vector-linked, encoded in the expression vector upstream of the cloning site of the target gene. As such, the lack of modularity of purification and solubility tags in expression vectors presents a bottleneck in high-throughput expression screens as the user is limited to the solubility tags encoded in the vectors available to them. Therefore, tackling the problem of modularity represents an opportunity to increase the efficiency of expression trials, and readily allows for the incorporation of novel purification and solubility tags as they are developed.

The pOPIN vector suite, generated by the Oxford Protein Production Facility (OPPF), is a set of expression vectors encoding various purification and solubility tags at either the N- or C-terminus of the gene of interest (GOI) (Berrow et al., 2007). These vectors allow for a straightforward cloning method via ligation independent cloning (LIC) and rapid generation of constructs. Furthermore, the pOPIN vectors are compatible with multiple hosts, with many being able to be used in bacterial, insect and mammalian cell hosts (Berrow et al., 2007). One shortcoming of these excellent vectors is a lack of modularity, meaning users are restricted to the solubility tags provided in the vector suite.

Golden Gate cloning (also known as Greengate cloning or MoClo) represents a fast and efficient method of cloning genes through the use of Type IIS restriction endonucleases that cut outside their recognition site to reveal user defined four nucleotide overhangs at both the 5’ and 3’ ends of the DNA (Engler et al., 2008). These overhangs can be exploited to allow scarless cloning, as well as for design and assembly of multiple DNA fragments in a single ligation reaction (Padgett and Sorge, 1996). They also allow for efficient subcloning between vectors (Engler et al., 2008). Golden Gate cloning allows for the generation of diverse “level 0” parts, which can be promoters, the GOI, terminators, epitope tags etc. These can then be assembled into “level 1” expression cassettes, which themselves can be further cloned along with other level 1 expression cassettes to give rise to “level 2” multi-gene assemblies. Moreover, the scarless nature of Golden Gate cloning makes the technique excellent for synthetic design approaches such as assembling chimeric proteins, allowing the generation of protein domain-swaps, or tagging proteins. Due to its high efficiency, modularity, and well established sequential cloning strategy, Golden Gate cloning has been incorporated in multiple vector systems for eukaryotic expression, such as plants or insect cells (Engler et al., 2014, 2009; Neuhold et al., 2020).

Here, we present a modified pOPIN vector suite we call pOPIN-GG that takes advantage of Golden Gate cloning (Engler et al., 2009, 2008) to incorporate modularity into the pOPIN vectors without disrupting efficacy of expression. These vectors also allow cross-compatibility with other Golden Gate systems, such as plant expression binary vectors (Engler et al., 2014; Patron et al., 2015), and simple one-pot reactions which allow for the bespoke construction of expression vectors containing the GOI and desired purification or solubility tags. We demonstrate that the pOPIN-GG vectors express comparable levels of protein to those of the classic pOPIN suite, whilst also enabling the user to selectively vary the choice of tags. This incorporation of modularity in protein expression supplied by pOPIN-GG vectors further advances high-throughput protein expression trials and subsequent preparative purification in *E. coli*.

## Results

### Cloning into the pOPIN-GG vectors

Cloning into the pOPIN-GG vectors follows standard Golden Gate protocols where matching overhangs between acceptor vector and inserts, revealed by digestion with a Type IIS endonuclease, *BsaI,* are ligated with T4 ligase (Engler et al., 2009, 2008). Here, we have engineered two sets of overhangs into the pOPIN-GG vectors: 5’ CCAT and 3’ GCTT for pOPIN-GG N-terminal tag compatible vectors (pPGN), and 5’ AATG and 3’ GCTT overhangs for pOPIN-GG C-terminal tag compatible vectors (pPGC). The pPGN and pPGC vectors have been developed with two different options for antibiotic selection, carbenicillin (the pPGN-C and pPGC-C vectors) and kanamycin (the pPGN-K and pPGC-K vectors), which allows for co-expression via bacterial co-transformation. Annotated vector maps of these pOPIN-GG acceptors are shown in Figure 1, detailing the changes made to the original pOPIN-F and pOPIN-A vectors (the modifications are further described in the experimental procedures).

**Figure 1.**
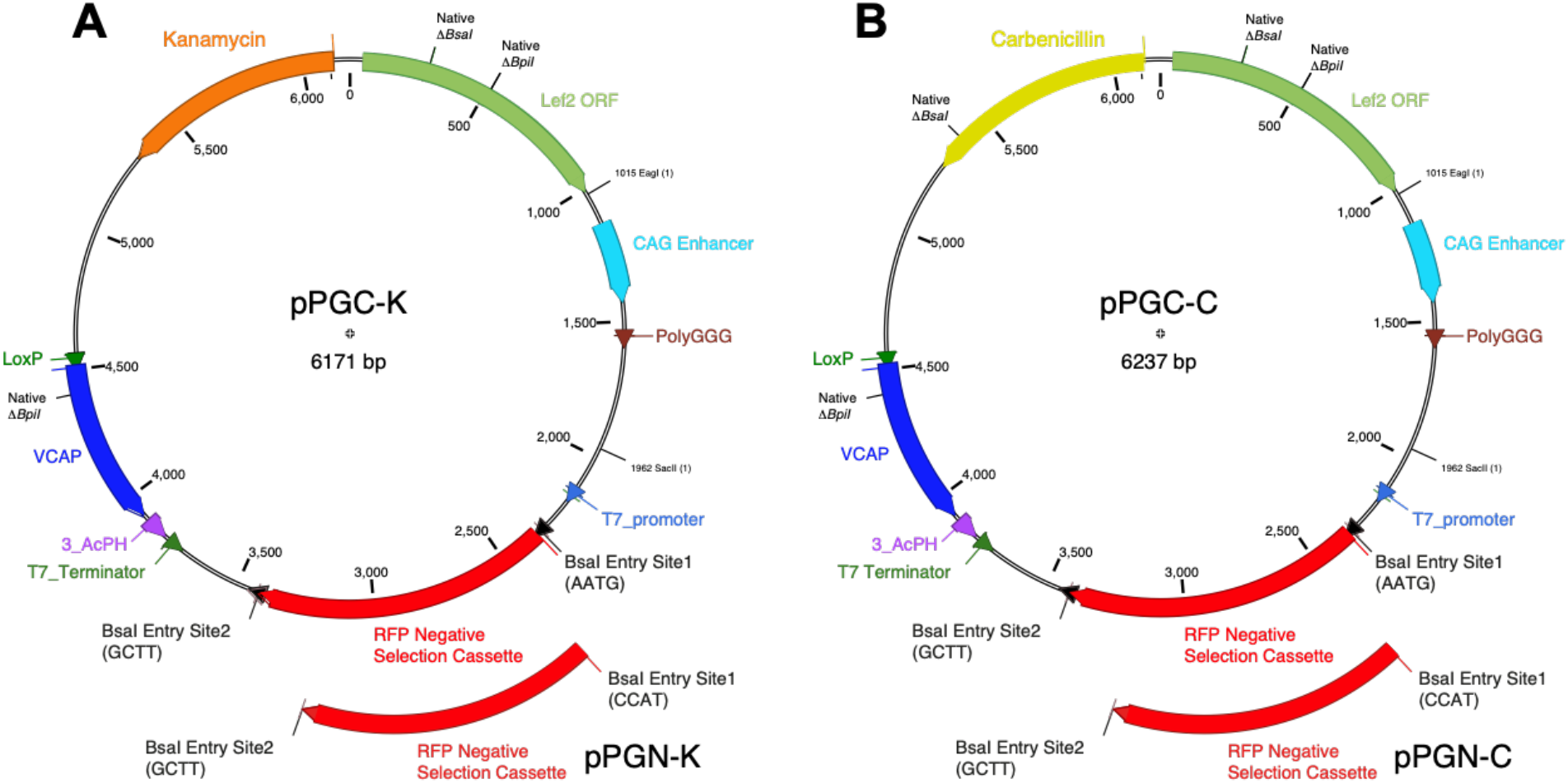
Annotated vector maps of the pOPIN-GG acceptors. pOPIN vectors pOPIN-A and pOPIN-F were modified to generate four pOPIN-GG acceptors. **A)** pPGN-K and pPGC-K are Kan*^R^* vectors descended from pOPIN-A for N-terminal and C-terminal tagging, respectively. **B)** pPGN-C and pPGC-C are Carb*^R^* vectors descended from pOPIN-F for N-terminal and C-terminal tagging, respectively. Δ*BsaI* and Δ*BpiI* represent domesticated *BsaI* and *BpiI* sites to allow for Golden Gate cloning compatibility. An RFP negative selection cassette was integrated into the backbone between the *BsaI* entry sites to allow for red/white selection upon insertion of the GOI.

Cloning of GOIs into the pOPIN-GG vectors are based on a common syntax for Type IIS endonuclease-mediated assembly (Patron et al., 2015). For cloning of N-terminally tagged GOIs into pPGN, the desired N-terminal tag must have the overhangs 5’ CCAT and 3’ AATG, and the GOI have the overhangs 5’ AATG and 3’ GCTT, when revealed by digestion with *BsaI.* For cloning into pPGC, the GOI requires the overhangs 5’ AATG and 3’ TTCG and the C-terminal tag must have the overhangs 5’ TTCG and 3’ GCTT. In addition, a GOI encoding an untagged protein can be cloned by using the 5’ AATG and 3’ GCTT overhangs and one of the C-terminal tagging compatible pPGC vectors, which contain the 5’AATG and 3’ GCTT overhangs. Figure 2 visualises the compatible overhangs between vector, tag and insert for cloning to produce N-terminally tagged, C-terminally tagged, and untagged proteins. Overhangs can be introduced into a desired sequence through either PCR with primers containing overhangs and a *BsaI* site (Table 1), or through *BsaI*-digestion of a compatible level 0 entry vector containing either tag or insert of interest. This highlights a major advantage of the pOPIN-GG vectors, by which a single PCR amplification or level 0 vector containing the GOI with the appropriate N-terminal or C-terminal tagging overhangs can be used for multiple simultaneous digestion-ligation reactions as the overhangs remain universal between acceptor, tag, and insert. pOPIN-GG acceptors along with multiple level 0 vectors containing N-terminal and C-terminal tags with the necessary overhangs for cloning are listed in Table 2.

**Table 1.**
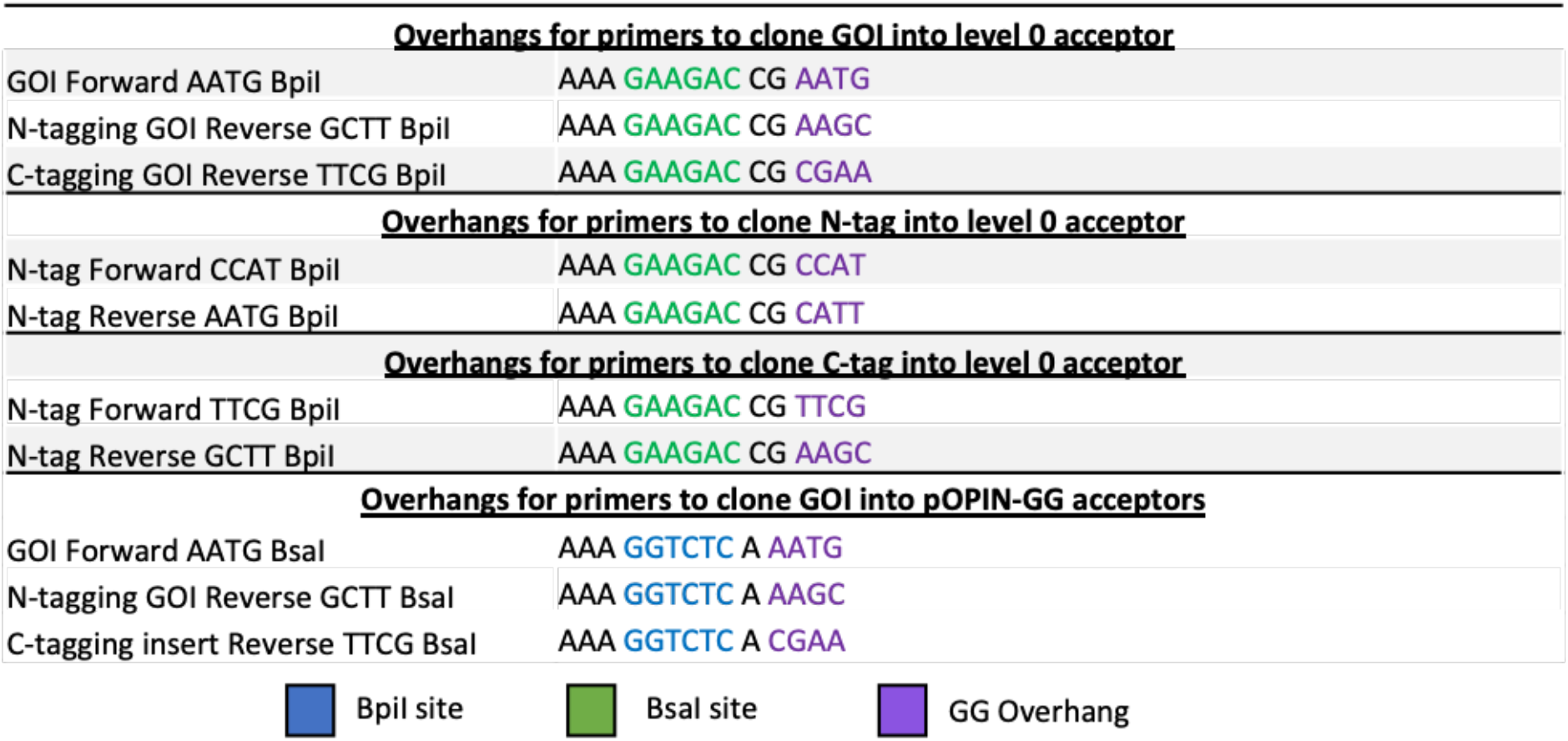
Overhang sequences for primers to clone tags and inserts into various acceptors.

**Table 2.** pOPIN-GG acceptors and compatible vectors available on Addgene.

| Vectors | Description | 5' Overhang | 3' Overhang | Selection | Addgene ID |
| --- | --- | --- | --- | --- | --- |
| **<u>Level 0 acceptors for making level 0 modules</u>** |  |  |  |  |  |
| plCH41308 | Acceptor for CDS to be N-tagged | AATG | GCTT | Spectinomycin | 174574 |
| plCSL01005 | Acceptor for CDS to be C-tagged | AATG | TTCG | Spectinomycin | 174575 |
| plCSL01002 | Acceptor for creating N-tag modules | CCAT | AATG | Spectinomycin | 174576 |
| plCSL01003 | Acceptor for creating C-tag modules | TTCG | GCTT | Spectinomycin | 174577 |
| **<u>Level 1 acceptors for protein expression</u>** |  |  |  |  |  |
| pPGN-C | Expression vector compatible with N-terminal tagging | CCAT | GCTT | Carbenicillin/Ampicillin | 174578 |
| pPGN-K | Expression vector compatible with N-terminal tagging | CCAT | GCTT | Kanamycin | 174579 |
| pPGC-C | Expression vector compatible with C-terminal tagging | AATG | GCTT | Carbenicillin/Ampicillin | 174580 |
| pPGC-K | Expression vector compatible with C-terminal tagging | AATG | GCTT | Kanamycin | 174581 |
| **<u>Premade level 0 modules encoding N-terminal tags</u>** |  |  |  |  |  |
| plCSL30015 | 6xHIS-MBP-3C | CCAT | AATG | Spectinomycin | 174582 |
| plCSL30017 | 6xHIS uncleavable | CCAT | AATG | Spectinomycin | 174583 |
| plCSL30018 | 6xHIS -SUMO-3C | CCAT | AATG | Spectinomycin | 174584 |
| plCSL30019 | 6xHIS-3C | CCAT | AATG | Spectinomycin | 174585 |
| plCSL30022 | Twin Streptavidin uncleavable | CCAT | AATG | Spectinomycin | 174586 |
| plCSL30028 | 6xHIS-GB1 | CCAT | AATG | Spectinomycin | 174587 |
| **<u>Premade level 0 modules encoding C-terminal tags</u>** |  |  |  |  |  |
| plCSL50001 | 6xHIS-TEV-3xFLAG | TTCG | GCTT | Spectinomycin | 174588 |
| plCSL50025 | 6xHIS uncleavable | TTCG | GCTT | Spectinomycin | 174589 |
| plCSL50026 | 3C-6xHIS | TTCG | GCTT | Spectinomycin | 174590 |
| plCSL50027 | Twin Streptavidin uncleavable | TTCG | GCTT | Spectinomycin | 174591 |

**Figure 2.**
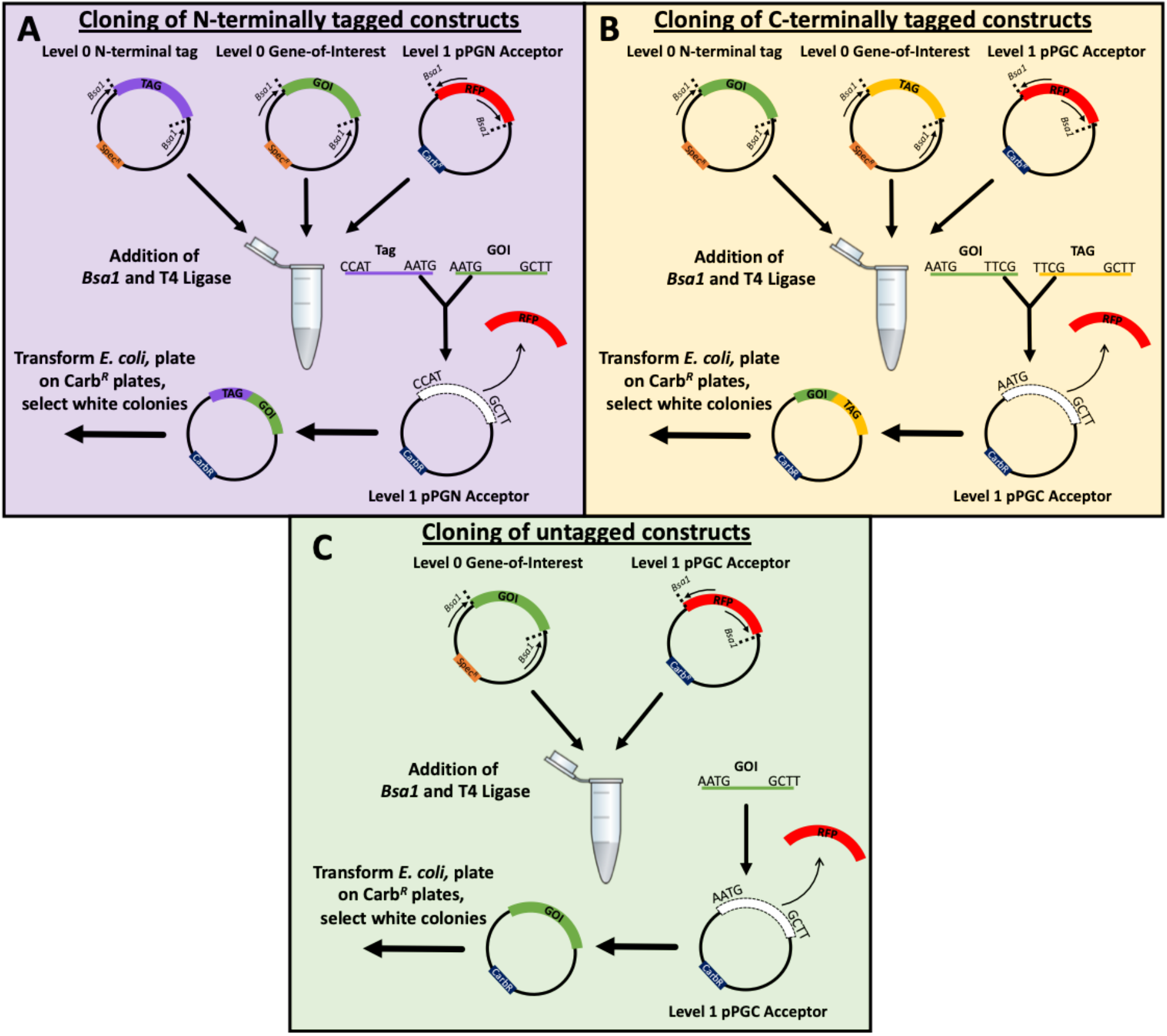
Cloning strategy flowcharts. The pOPIN-GG vectors enable N-terminally tagged, C-terminally tagged, or untagged constructs. **A)** Cloning strategy for N-terminal tagging. GOI requires overhangs 5’ AATG and 3’ GCTT which can be revealed from *BsaI* treatment of a level 0 vector or added by PCR. The GOI can then been combined with one of the pPGN acceptor and N-terminal tag level 0 vectors to generate an N-terminally tagged level 1 construct. **B)** Cloning strategy for C-terminal tagging. GOI requires overhangs 5’ AATG and 3’ TTCG which can be revealed from *BsaI* treatment of a level 0 vector or added via PCR. The GOI can then be combined with one of the pPGC acceptor and C-terminal tag level 0 vectors to generate a C-terminally tagged level 1 construct. **C)** Cloning strategy for generating an untagged gene. As for N-terminal tagging, the GOI requires overhangs 5’ AATG and 3’ GCTT which can be revealed from *BsaI* treatment of a level 0 vector or added by PCR. However, is then combined with one of the pPGC vectors, resulting in an untagged GOI, useful in co-expression. Carb*^R^* level 1 acceptor vectors are shown for simplicity, Kan*^R^* level 1 acceptor vectors can be used instead as required.

### The pOPIN-GG vectors maintain protein expression levels comparable to the original pOPIN vectors

To test the efficacy of the pOPIN-GG vectors for protein expression, we cloned AVR-PikF (a small, secreted effector protein from the blast fungus *Magnaporthe oryzae* (Longya et al., 2019)), into the vectors pOPIN-F (N-terminal 6xHIS-3C), pOPIN-S3C (N-terminal 6xHIS-SUMO-3C), pOPIN-M (N-terminal 6xHIS-MBP-3C) and pOPIN-E (C-terminal 6xHIS uncleavable), and generated the equivalent constructs using our pOPIN-GG system (see experimental procedures). We also cloned AVR-PikF with a 6xHIS-GB1 solubility tag (Kobashigawa et al., 2009), to which we did not have access to the pOPIN equivalent, to demonstrate the adaptability of the pOPIN-GG system. Figure 3 shows a comparison between AVR-PikF expressed in the *E. coli* SHuffle strain using the gene cloned in the original pOPIN vectors and the pOPIN-GG vectors and after benchtop Ni^2+^-immobilised metal affinity chromatography (IMAC) purification. We observed no significant differences in the yield of protein between the pOPIN vectors and new pOPIN-GG vectors, demonstrating the pOPIN-GG vectors retain the same capacity for protein production as the parent vectors.

**Figure 3.**
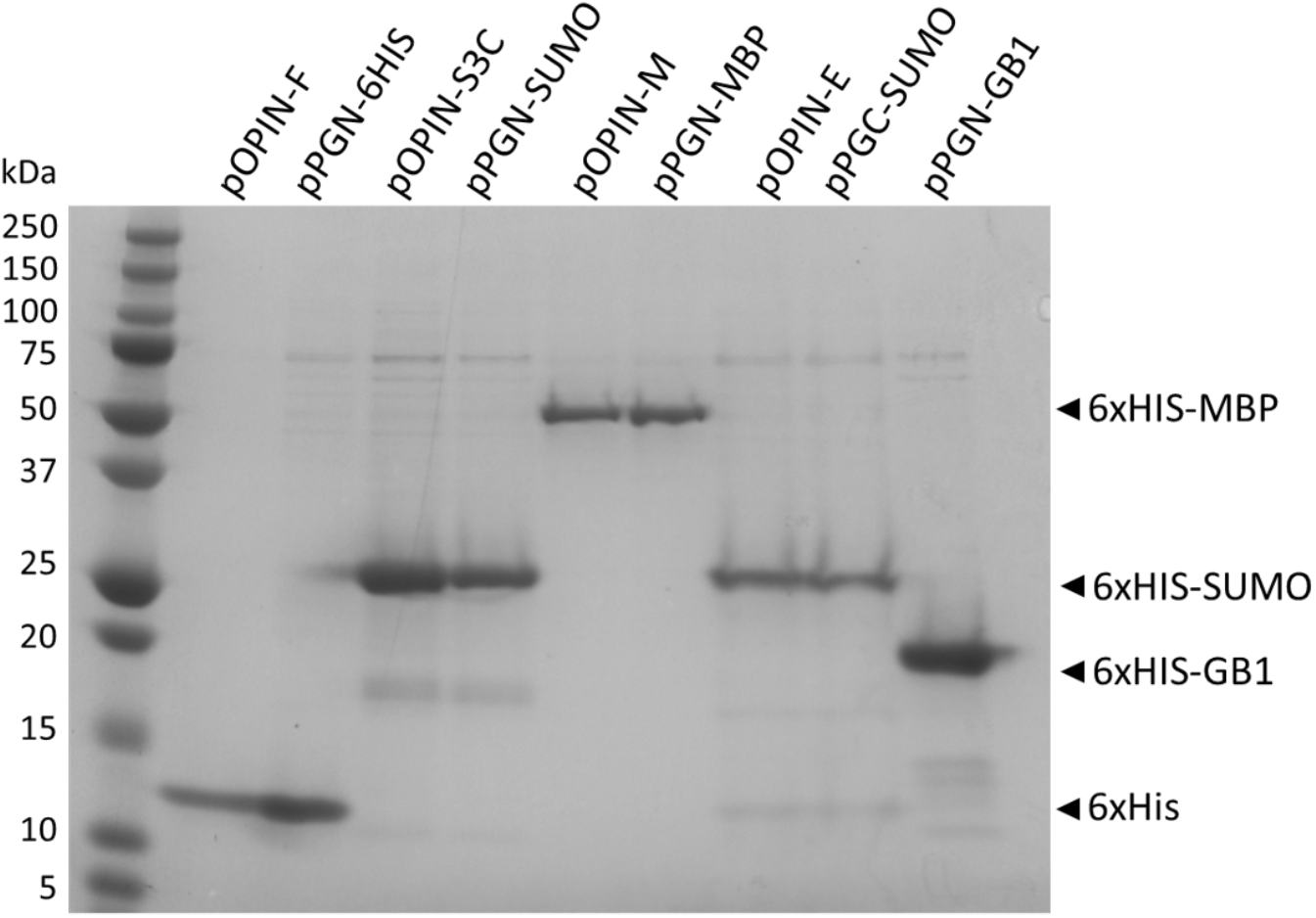
Expression and purification of AVR-PikF in pOPIN vectors vs AVR-PikF in pOPIN-GG vectors. AVR-PikF was cloned into the pOPIN vectors pOPIN-F, pOPIN-S3C, pOPIN-M, and pOPIN-E (with additional SUMO tag to aid solubility). The equivalent constructs were produced in the pOPIN-GG system, as well as an additional 6xHIS-GB1-tagged construct. Constructs were expressed in *E. coli* SHuffle cells and purified using Ni-NTA affinity resin before being visualised by SDS-PAGE.

### The pOPIN-GG vectors are compatible with co-expression

To further demonstrate the flexibility of the pOPIN-GG vectors, we next cloned an interaction partner of AVR-PikF, OsHIPP19 (Maidment et al., 2021), into pPGN-C with a 6xHIS-GB1 tag. We co-transformed *E. coli* SHuffle cells with the 6xHIS-GB1-OsHIPP19 vector and an untagged variant of AVR-PikF (in pPGC-K), expressed the proteins, and performed Ni^2+^-IMAC coupled with size exclusion chromatography (SEC) to purify a complex between AVR-PikF and OsHIPP19 (Figure 4). SDS-PAGE shows both proteins are expressed and purified, with AVR-PikF co-purified along with OsHIPP19 (Figure 4). These results demonstrate the pOPIN-GG vectors can be used for efficient expression and purification of protein complexes.

**Figure 4.**
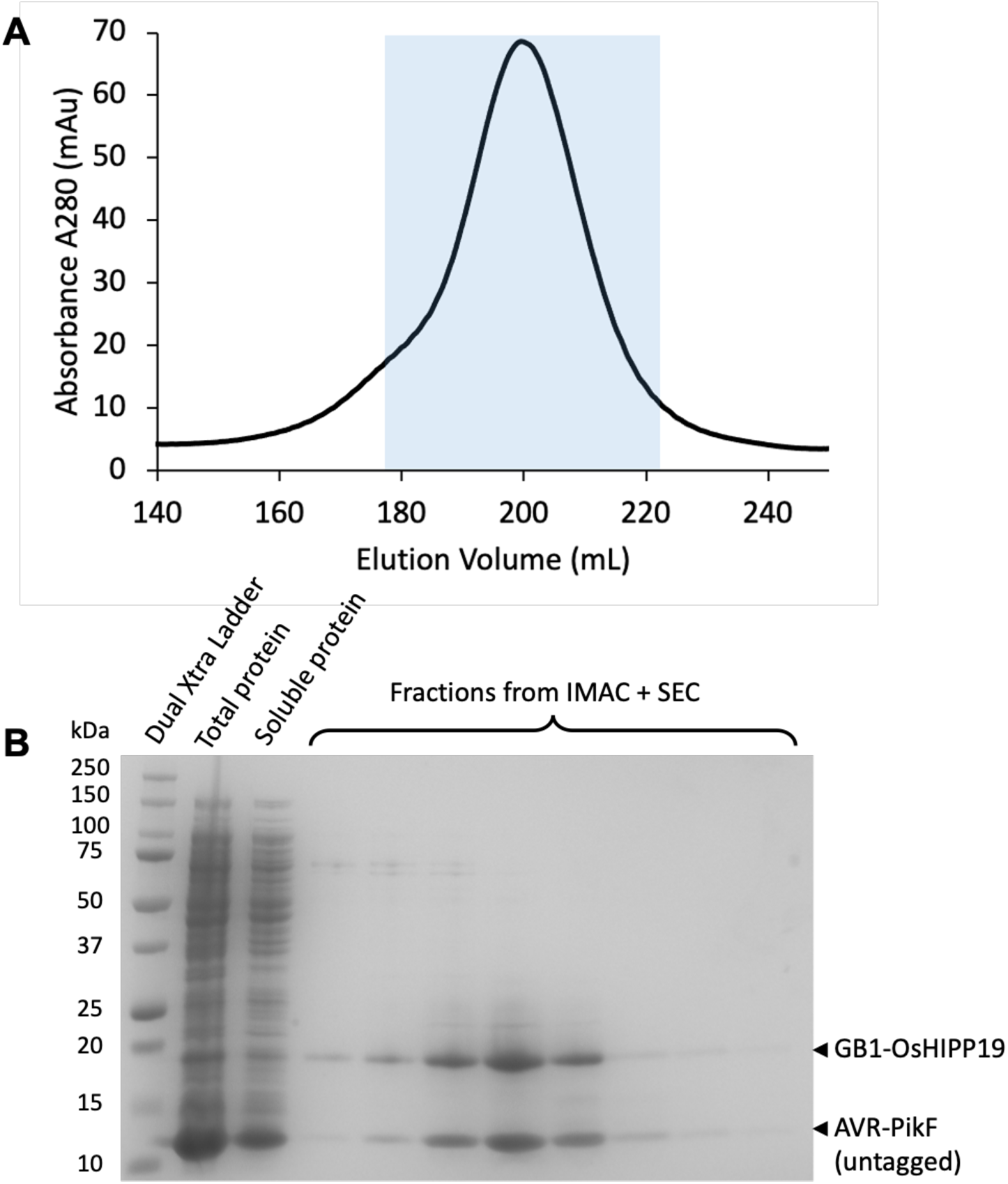
Co-expression and purification of AVR-PikF and OsHIPP19 using the pOPIN-GG system. An untagged construct of AVR-PikF was generated using pPGC-K and co-expressed with OsHIPP19 cloned with a 6xHIS-GB1 solubility tag in pPGN-C *E. coli* SHuffle cells. Constructs were purified using Ni^2+^-mediated immobilised metal affinity chromatography (IMAC) coupled with size exclusion chromatography (SEC) before visualised via SDS-PAGE. **A)** SEC chromatogram of the AVR-PikF / OsHIPP19 complex. Blue box indicates the fractions analysed by SDS-PAGE. **B)** Instant blue^®^ Coomassie-stained SDS-PAGE gel of the fractions from SEC showing the co-elution of the AVR-PikF and OsHIPP19 proteins, indicating complex formation.

## Discussion

The ability to test multiple purification and solubility tags in *E. coli* expression trials is important for determining the best conditions for protein production. High-throughput expression trials allow the user to explore potential avenues for successful protein production, and a key step is the ability to clone the gene of interest easily and effectively into a variety of constructs with different purification and solubility tags. Here we describe pOPIN-GG, a modification of the pOPIN vector suite, which allows for modular cloning using Golden Gate. The redesigned pOPIN-GG vector suite retains the expression levels of the pOPIN vectors (with the proteins we tested) while enhancing their modularity. A major advantage of the pOPIN-GG system is it allows for the rapid introduction new purification and solubility tags as they emerge, in a straightforward and easy to clone manner. The different pOPIN-GG acceptors are cross-compatible for co-expression, allowing for ease of co-expression and purification of protein complexes. Finally, the utility of the pOPIN-GG vectors presented here have already been successfully tested by the community for the expression and purification of a fungal effector from *Parastagonospora nodorum* (Outram et al., 2021), as well as for detailing the structural mechanisms underpinning the evolution of *Magnaporthe oryzae* effectors for a specific host target (Bentham et al., 2021).

## Conclusions

The pOPIN vector suite is a well-established and effective *E. coli* vector system for protein expression and purification. By modifying the pOPIN vectors to be Golden Gate compatible we have introduced modularity to the system, providing the user with the advantages of modular cloning and allowing the incorporation new purification and solubility tags to the vector repertoire, streamlining the vector assembly step, with no cost to the expression efficacy of the system. We have made the pOPIN-GG vectors available on Addgene (https://www.addgene.org/Mark_Banfield) as a package with level 0 and level 1 acceptors, and level 0 N-terminal and C-terminal tag constructs.

## Experimental Procedures

### Domestication and modification of the pOPIN vectors to generate the pOPIN-GG vectors

We used the pOPIN-F vector (Berrow et al., 2007) as template to generate Golden Gate compatible versions of the pOPIN vectors, making considerable effort to minimise alterations to the existing pOPIN-F vector backbone. To adapt pOPIN-F to the Golden Gate cloning system, we removed all of the native Type IIS *BsaI* and *BpiI* restriction sites from the sequence, a process traditionally termed ‘domestication’. After domestication, we generated two different antibiotic resistance variants (carbenicillin and kanamycin) conducive to the need for dual protein co-expression. We then took the carbenicillin resistant (Carb*^R^*) and kanamycin resistant (Kan*^R^*) vectors and engineered two variants for each to allow for the insertion GOIs compatible with either N-terminal or C-terminal tags. To do this, we reintroduced two *BsaI* sites between the T7 promoter and terminator, which would yield 5’ CCAT and 3’ GCTT nucleotide overhangs for the N-terminal tag compatible vectors, and 5’ AATG and 3’ GCTT overhangs in the C-terminal tag compatible vectors post treatment with *BsaI* (Figure 1). Ultimately, we developed four pOPIN-GG acceptor vectors, pPGN-C (Carb*^R^*) and pPGN-K (Kan*^R^*) for N-terminal tagging, and pPGC-C (Carb*^R^*) and pPGC-K (Kan*^R^*) for C-terminal tagging (Table 1). Further, to assist with positive clone identification, we introduced a visible red fluorescent protein (RFP) negative selection marker, allowing users to select positive white colonies after transformation. Importantly, the Carb*^R^* and Kan*^R^* versions of the acceptors also contain different origins of replication to allow for efficient co-expression in addition to co-transformation.

To overcome complications with PCR (polymerase read-through in the native pOPIN-F vector), we removed a 15xG homopolymer string by excising a 947bp (*EagI-SacII*) section of vector backbone using traditional restriction enzymes and stored this for later use. As a consequence of increased Golden Gate assembly efficiency from cloned (i.e. non-linear) fragments, the remaining sections of the backbone (incorporating changes to the native 2x*BsaI* and 2x*BpiI* restriction sites) were amplified by PCR and cloned into custom level 0 acceptors. Once independently sub-cloned, the mutated fragments could subsequently be re-assembled (along with the *EagI-SacII* cassette), eliminating all the existing *BsaI* and *BpiI* restriction sites to generate a fully domesticated version of the original pOPIN-F vector.

Following domestication, larger sections of the vector back-bone could be re-cloned into additional custom level 0 acceptors to enable a pre-domesticated iGEM RFP negative selection reporter cassette (originating from *Discosoma striata*) to be inserted in place of the pOPIN-F N-terminal 6xHIS tag and Lac operon (LacZ) negative selection elements. The border sequences immediately flanking this region were found to be important for protein expression and were modified to directly mimic those of the native pOPIN vectors. Final vector assembly was achieved by ligating vector back-bone fragments together along with the *EagI-SacII* and RFP negative selection reporter elements. Changes to the Golden Gate cloning insert overhangs, and exchange of the vector bacterial antibiotic resistance, were achieved by making the necessary modifications to the respective level 0 assembly components.

### Cloning of AVR-PikF into pOPIN and pOPIN-GG vectors for test expression and purification

The AVR-PikF coding sequence (encoding residues starting from the end of the signal peptide 21 – 113) was cloned into pOPIN and pOPIN-GG vectors via Infusion cloning and Golden Gate, respectively. Infusion cloning of AVR-PikF in the pOPIN vectors, pOPIN-F, pOPIN-S3C, pOPIN-M and pOPIN-E was performed as described by (Berrow et al., 2007). To clone AVR-PikF into the pOPIN-GG constructs, the AVR-PikF sequence was amplified with primers that introduced overhangs at the 5’ and 3’ of the sequence containing a *BsaI* Type IIS endonuclease sites (Table 1) which would reveal 5’ CCAT and 3’ GCTT overhangs or 5’ AATG and 3’ TTCG overhangs, for N-terminal tagged or C-terminal tagged constructs, respectively (Figure 1). AVR-PikF amplicons were subsequently used in a one-pot Golden Gate reaction (Engler et al., 2009, 2008) with the pPGN-C acceptor for N-terminal tagging or the pPGC-C for C-terminal tagging and desired N-/C-terminal tag-containing construct (Figure 1). Golden Gate reactions were then transformed into chemicompetent Stellar *E. coli* cells (Takara Bio) and positive clones were identified via RFP selection before sequencing to confirm the cloning was successful.

### Small Scale Expression and Purification of differentially tagged AVR-PikF in pOPIN and pOPIN-GG vectors

Constructs for protein expression were transformed into *E. coli* SHuffle cells (NEB) via heat shock. Protein expression was performed via autoinduction (Studier, 2005) under the required selection. Cells were harvested by centrifugation and resuspended in lysis buffer (50 mM HEPES pH 7.4, 500 mM NaCl, 30 mM imidazole, 5 mM glycine, 5% glycerol) at a ratio of 5:1 buffer per gram of cell pellet. 8 mL of resuspended cells were transferred to a 24 well plate and lysed by sonication with a 24-tip sonication horn. Cell lysate was clarified by centrifugation at 45,000 *g* for 20 mins. 500 μl of Ni-NTA resin (Qiagen) was added to each of the clarified cell lysates and incubated for 20 mins with gentle shaking. After incubation, cell lysates with Ni-NTA resin were transferred to 15 ml gravity flow columns and Ni-NTA resin was washed with 2 column volumes of lysis buffer to remove non-specific interactors. Proteins were eluted from the Ni-NTA resin with 1 mL of elution buffer (50 mM HEPES pH 7.4, 500 mM NaCl, 500 mM imidazole). Samples were visualised by SDS-PAGE with 16 % Teo-tricine polyacrylamide gels (Abcam) stained with Instant Blue^®^ Coomassie Stain (Abcam).

### Expression and purification of the AVR-PikF and OsHIPP19 complex

A 6xHIS-GB1-tagged OsHIPP19 in pPGN-C was co-transformed with an untagged AVR-PikF in pPGC-K into *E. coli* SHuffle cells and plated on carbenicillin + kanamycin LB agar plates. Protein expression was performed via autoinduction (Studier, 2005) under carbenicillin + kanamycin selection. Cells were harvested by centrifugation and resuspended in lysis buffer (50 mM HEPES pH 7.4, 500 mM NaCl, 30 mM imidazole, 5 mM glycine, 5% glycerol) at a ratio of 5:1 buffer per gram of cell pellet. Cells were lysed via sonication and lysate was clarified by centrifugation at 45,000 *g*. Clarified lysate was then subjected to Nickel immobilised metal affinity chromatography (IMAC) followed by size exclusion chromatography (SEC) with a Superdex S75 26/600. Fractions from SEC were visualised by SDS-PAGE with 16 % Teo-tricine polyacrylamide gels (Abcam) stained with Instant Blue^®^ Coomassie Stain (Abcam).

## Acknowledgements

We thank Ray Owens from the Oxford Protein Production Facility for provision of the pOPIN vectors. We thank Dr. Florian Altegoer for kindly providing pEMGB1 vector for the cloning of 6xHIS-GB1 tag. This work was supported by UKRI-BBSRC (grant BB/P012574), UKRI-BBRSC Norwich Research Park Biosciences Doctoral Training Partnership, (grant BB/M011216/1, for FAV) and the ERC (proposal 743165), the ERAMUS^+^ training programme, the John Innes Foundation and the Gatsby Charitable Foundation.

## Data Availability Statement

All data supporting the findings of this study are contained within the manuscript. The pOPIN-GG vectors are available directly through Addgene (https://www.addgene.org/Mark_Banfield/).

